# Watching antibiotics in action: Exploiting time-lapse microfluidic microscopy as a tool for target-drug interaction studies in *Mycobacterium*

**DOI:** 10.1101/477141

**Authors:** Damian Trojanowski, Marta Kołodziej, Joanna Hołówka, Rolf Müller, Jolanta Zakrzewska-Czerwińska

## Abstract

Spreading resistance to antibiotics and the emergence of multidrug-resistant strains have become frequent in many bacterial species, including mycobacteria. The genus *Mycobacterium* encompasses both human and animal pathogens that cause severe diseases and have profound impacts on global health and the world economy. Here, we used a novel system of microfluidics, fluorescence microscopy and target-tagged fluorescent reporter strains of *M*. *smegmatis* to perform real-time monitoring of replisome and chromosome dynamics following the addition of replication-altering drugs (novobiocin, nalidixic acid and griselimycin) at the single-cell level. We found that novobiocin stalled replication forks and caused relaxation of the nucleoid, nalidixic acid triggered rapid replisome collapse and compaction of the nucleoid, and griselimycin caused replisome instability with subsequent over-initiation of chromosome replication and over-relaxation of the nucleoid. This work is an example of using a microscopy-based approach to evaluate the activity of potential replication inhibitors and provides mechanistic insights into their modes of action. Our system also enabled us to observe how the tested antibiotics affected the physiology of mycobacterial cells (i.e., growth, chromosome segregation, etc.). Because proteins involved in the DNA replication are well conserved among bacteria (including mycobacterial species), the properties of various replication inhibitors observed here in fast-growing *M. smegmatis* may be easily extrapolated to slow-growing pathogenic tubercle bacilli, such as *M. tuberculosis*.

**Significance:** The growing problem of bacterial resistance to antibiotics and the emergence of new strains that are resistant to multiple drugs raise the need to explore new antibiotics and re-evaluate the existing options. Here, we present a system that allows the action of antibiotics to be monitored at the single-cell level. Such studies are important in the light of bacterial heterogeneity, which may be enhanced in unfavorable conditions, such as under antibiotic treatment. Moreover, our studies provide mechanistic insights into the action modes of the tested compounds. As combined therapies have recently gained increased interest, it is also notable that our described system may help researchers identify the best combination of antimicrobials for use against infections caused by a variety of bacteria.

## Introduction

Bacterial resistance to antibiotics, which is an increasing health problem worldwide, is a concern for every commercially used antimicrobial (1–7). Recent years have seen the constant emergence of new strains that are resistant to multiple drugs (multidrug-resistant, MDR, and extensively resistant, XDR), including last-resort antibiotics (e.g., strains of *Neisseria gonorrhoeae* that resist third-generation cephalosporins, strains of *Klebsiella pneumoniae* that resist carbapenems and strains of *Enterobacteriaceae* that resist colistin) (8). Moreover, relatively few novel compounds have been approved for managing bacterial infections in recent years, with fewer than a dozen such drugs approved in the last 10 years. Therefore, novel antibiotics are urgently needed.

Antibiotics target basic cellular processes, such as the synthesis and integrity of the cell wall (penicillins, cephalosporins, lipoglycopeptides, polymyxins, etc.), transcription (rifampicin), translation (aminoglycosides, macrolides, linkosamides, tetracyclines, oxazolidinones), and metabolic pathways (sulphonamides, diaminopyrimidines) (9–13). Since the proteins that govern bacterial DNA replication differ from their eukaryotic counterparts, chromosome replication represents another promising therapeutic target (14–17). However, bacterial chromosome replication is targeted by only a few current antibiotics, such as quinolones, aminocoumarins and metronidazole (18–23). Recently, it has also been reported that nonsteroidal anti-inflammatory drugs exert an inhibitory effect on bacterial chromosome replication (24). Interestingly, quinolones, which are important anti-tuberculosis (TB) drugs, are among the most frequently prescribed antibiotics in modern medicine.

The genus *Mycobacterium* encompasses both human (*M. leprae* and *M. tuberculosis*) and animal (*M. bovis*) pathogens that cause severe diseases and have profound impacts on global health and the world economy. Although the number of new *M. tuberculosis* (*Mtb*) infection cases has been decreasing annually (25), TB remains one of the most prominent causes of death worldwide and the main cause of death among HIV-infected individuals (26). It is estimated that one-third of the human population is latently infected with *Mtb* and that tubercle bacilli may be reactivated from the latent state upon immunosuppression later in life (27). As with other pathogens, resistance of *Mtb* is becoming a serious obstacle in effective drug therapy. According to the WHO, of the 10 million new TB cases in 2016, nearly half a million were classified as MDR-TB (resistant to two anti-TB drugs), and among them, about 6% were caused by XDR strains (resistant to more than four anti-TB medications)(26). The increasing number of resistant *Mtb* strains coupled with the short list of anti-TB drugs prompted researchers to reevaluate some commonly used antibiotics (e.g., linezolid, clofazimine, amoxicillin/clavulanate) for off-label treatment of TB (28, 29). One serious challenge in the treatment of TB caused by both susceptible and resistant *Mtb* strains is the high population heterogeneity of mycobacterial cells (30–38). Asymmetric growth (mycobacteria elongate preferentially from the old pole) and asymmetric septum placement give rise to daughter cells of unequal sizes and growth rates and, as indicated by some reports, different susceptibilities to antibiotics (30, 39, 40). Moreover, exposure to stress further diversifies the population (41, 42), suggesting that mycobacteria utilize heterogeneity as key survival strategy under stressful conditions. Thus, there is a critical need for researchers to explore how anti-TB drugs act on individual cells.

Single-cell techniques have an advantage over traditional bath cultures in terms of providing insights into the mechanism of action of tested compounds, assuming that an appropriate reporter system is available (strains carrying fluorescent fusions to drug-targeted proteins, reporter genes, metabolic pathway indicators, etc.). However, only a few studies have examined the direct impact of antibiotics at single-cell resolution. Such studies are of great interest given that bacterial heterogeneity is seen under non-optimal conditions in the host environment and is likely to contribute to antimicrobial tolerance and/or resistance.

Here, we present a system that combines time-lapse microfluidic microscopy (TLMM) and replisome-tagged (Fig. 1A) fluorescent strains of *M. smegmatis* to allow real-time observation of how antibiotics affect chromosome replication. We show how the replisome and chromosome dynamics is altered upon the addition of novobiocin (an aminocoumarin), nalidixic acid (a quinolone) and griselimycin (43) (a novel antimicrobial agent), all of which exhibit different modes of action. To date, these antibiotics have been analyzed entirely using *in vitro* or batch studies. Because the proteins involved in DNA replication are highly conserved among mycobacteria, our results regarding the properties of various replication inhibitors obtained in fast-growing *M. smegmatis* may be easily extrapolated to the slow-growing tubercle bacilli, such as *M. tuberculosis*.

**Fig. 1.**
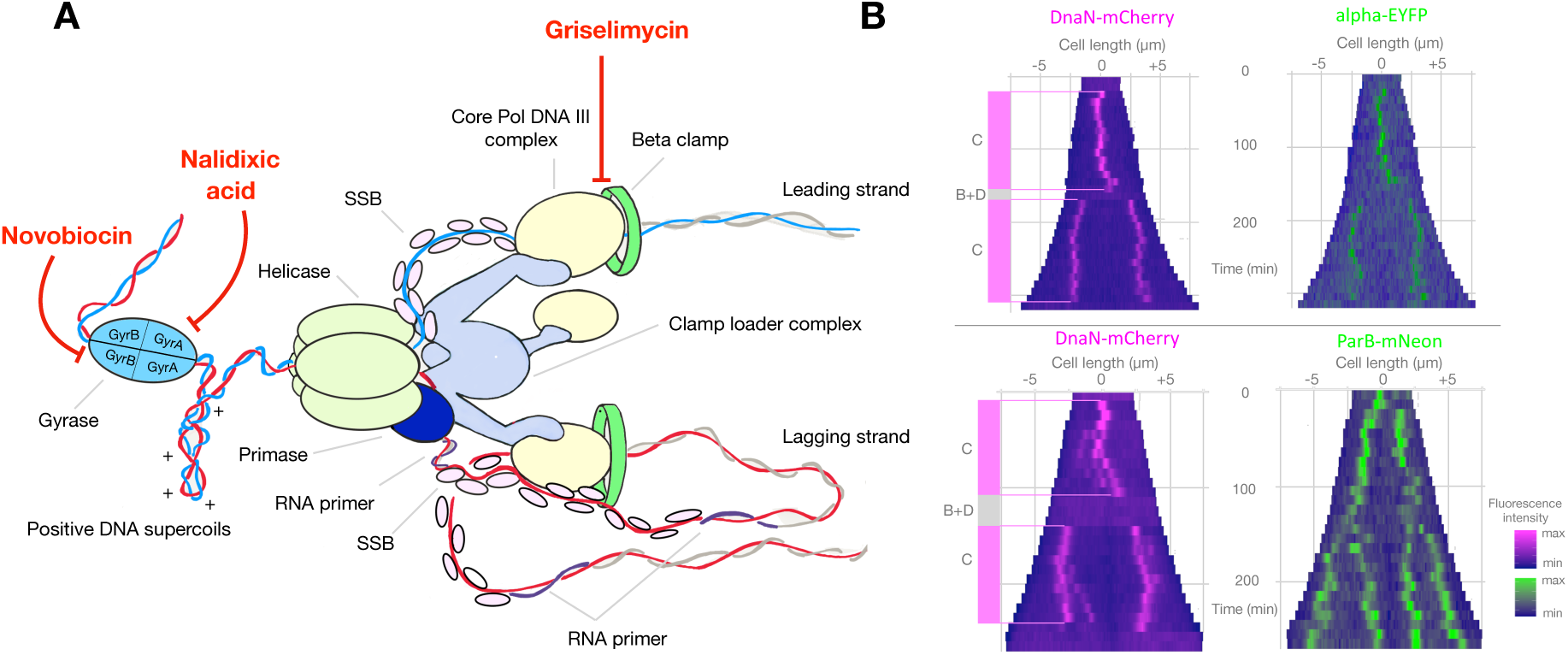
Replisomes are highly dynamic entities in *M. smegmatis*. (A) A schematic representation of the replisome, which is a multiprotein complex engaged in DNA replication. Core DNA polymerase III (core DNA Pol III) is loaded by the clamp loader complex via the tau subunit, whereas the beta clamp is loaded via the delta subunit of the clamp loader complex. The catalytic subunit alpha of core DNA Pol III interacts with the beta clamp in its hydrophobic cleft to give the replisome a high degree of processivity. Both Ndx and novobiocin target DNA gyrase. Novobiocin binds to the GyrB subunit, while Ndx binds to the gyrase/DNA complex (cleavable complex). Griselimycin abolishes the alpha-beta interactions by binding in the hydrophobic cleft of the sliding clamp. (B) Kymographs of representative cells from two *M. smegmatis* strains: DnaN-mCherry/Alpha-EYFP (top panel) and DnaN-mCherry/ParB-mNeon (bottom panel). The fluorescence intensities over time are depicted in magenta for DnaN-mCherry and in green for Alpha-EYFP or ParB-mNeon. “C period” refers to the time during which DnaN-mCherry and/or Alpha-EYFP is/are observed as diffraction-limited foci, while “B+D period” refers to the time between the termination of DNA replication and the initiation of another round of replication in the next generation.

As expected given that the tested antibiotics had different modes of action, we observed different cellular responses during the antibiotic treatments, particularly in terms of the replisome and chromosome dynamics. We thus describe a microscopy-based approach that can be used to evaluate the activity of potential replication inhibitors, while also providing mechanistic insights into the action modes of the studied drugs. The system described herein can allow researchers to simultaneously observe the target along with other processes (e.g., replication, growth and segregation), and thus provides additional results beyond the simple measurement of target protein inhibition.

## Results

### TLMM allows changes in chromosome and replisome dynamics to be observed in real-time during antibiotic treatment

In this study, we used fluorescent reporter strains of *M. smegmatis* and a microfluidic CellASIC Onix platform to observe in real-time the actions of novobiocin, nalidixic acid and griselimycin at the single-cell level. The bacterial replisome (a multiprotein complex that is involved in chromosome replication) and the targets of the studied antibiotics are schematically depicted in Figure 1A. Both nalidixic acid (Ndx) and novobiocin affect replisome passage indirectly by inhibiting the enzymatic activity of DNA gyrase, which normally triggers relaxation of positive supercoils ahead of the replication fork to resolve the torsional tension and allow DNA synthesis to proceed. Although the two drugs act on the same target, Ndx prevents the re-ligation of cleaved DNA by binding to the gyrase/DNA cleavable complex, resulting in double-strand breaks (44, 45), whereas novobiocin competes with ATP for binding to the GyrB subunit, and thus inhibits cleavage but not the binding of DNA (46–48). Ndx (as well as other quinolones) also inhibits the activity of topoisomerase IV (another type-II topoisomerase), which is crucial for resolving decatenates and chromosome dimers. More importantly, *M. tuberculosis* possesses only one type-II topoisomerase (i.e., DNA gyrase) whose activity combines those of a classical DNA gyrase (found in other bacteria) and topoisomerase IV (49, 50). In contrast to novobiocin and Ndx, griselimycin acts directly on the replication machinery; it prevents the interaction between the beta-clamp and the catalytic subunit alpha of DNA polymerase III to hinder the processivity of the replisome (43). We used previously constructed strains in which replisome subunits (catalytic subunit alpha and/or the beta-clamp), a chromosomal marker (HupB) and/or *oriC* (ParB bound to *oriC*-proximal *parS* sites) were tagged with different fluorescent proteins (FPs) (51–54). The strains used in this study are presented in Table S1. The growth of all strains was similar to that of the wild-type (WT) *M. smegmatis* mc^2^155 strain (Fig. S1 bottom right panel). We calculated the inhibitory concentration (IC) of the three tested antibiotics for all fluorescent reporter strains using a Bioscreen C instrument (see Methods and Fig. S2). The concentrations at which growth was inhibited by 50% (IC_50_) are presented in Table 1. For novobiocin and griselimycin, the IC_50_ values were lower for the reporter strains compared to WT cells, indicating that the reporter strains had higher susceptibilities to the tested antibiotics. This emphasizes the importance of determining inhibitory values for any strain selected for study.

**Table 1.** IC50 concentration of tested antibiotics for various strains used in the study.

| Strain | Antibiotic / IC50 (µg/ml) |  |  |
| --- | --- | --- | --- |
|  | Novobiocin | Nalidixic acid | Griselimycin |
| mc<sup>2</sup> 155 | 4.8 | 50 | 0.43 |
| DnaN-mCherry / ParB-mNeon | 4.0 | 50 | 0.22 |
| DnaN-mCherry / α-YFP | 1.2 | 50 | 0.12 |
| DnaN-mCherry / HupB-EGFP | 2.9 | 49 | 0.17 |

To examine replisome and chromosome dynamics in cells treated with sublethal doses of the selected antibiotics, we used concentrations of 5x and 10x IC_50_, which were expected to trigger discrete and observable changes without rapidly killing the bacterial cells. Cells loaded into microfluidic chambers were observed using the same protocol: 5 hours of growth under optimal conditions followed by 5 hours of antibiotic treatment (this constituted approximately twice the chromosome replication time) and 7 hours of washout. In the absence of any antibiotic, cells usually initiated one replication round per cell cycle (excluding the 10-15% multifork cells observed herein, which was consistent with a previous studies (51, 52)). Replication initiation corresponded to the appearance of a fluorescent spot that was positioned slightly asymmetric to the mid-cell. Under our experimental conditions, replication lasted for 119 +/− 16 min (C period, n=60) in the DnaN-mCherry/ParB-mNeon strain and 149 +/− 9 min (n=64) in the DnaN-mCherry/α-EYFP strain; thereafter, replication was terminated, which was observed as the disappearance of the DnaN-mCherry and/or α-YFP signal(s). This was followed by a period during which the fluorescence signal of the tagged replisomes was dispersed (B+D period); it lasted 27 +/− 10 min (n=60) in DnaN-mCherry/ParB-mNeon strains and 13 +/−11 (n=81) minutes in DnaN-mCherry/α-YFP strains (see kymographs in Fig. 1B). Contrary to the recent observations (55) in *E. coli* and *B. subtilis,* but consistent with our previous findings (51, 52), in *M. smegmatis*, we observed that replisomes frequently split and merged back together during C period.

### Novobiocin stalls replication forks but only moderately affects the cell elongation rate

In the presence of novobiocin, the replisomes (both Alpha-EYFP and DnaN-mCherry) remained visible but exhibited significantly decreased mobility along the cell (Fig. 2A-C, Supp. Movie 1). Because Alpha-EYFP fusion causes prolongation of the C period and shortening of the B+D period ((52) and see kymographs in Fig. 1B), we used DnaN-mCherry/ParB-mNeon to analyze the changes in replisome dynamics during novobiocin treatment. After the addition of novobiocin, the mobility of replisomes decreased in a dose-dependent fashion (Fig. 2C). As a consequence of this replication fork hold-up, the C period was profoundly prolonged under both 5x IC_50_ and 10x IC_50_ novobiocin (mean 178% and 195%, respectively, when we compared the mean C period in cells that terminated replication during novobiocin treatment versus that in untreated cells; n=50 cells per group). At a dose of 25x IC_50_ novobiocin, the replisomes were almost completely stalled. Two major group of cells were observed during novobiocin exposure, as depicted in the representative kymographs presented in Figure 2A and B. The first group (45% of 92 cells in the 5x IC_50_ group and 34% of 187 cells in the 10x IC_50_ group) comprised cells in which replisome foci were visible throughout the antibiotic treatment due to the delay in replication fork passage (black arrows on the kymographs). The second group comprised cells that terminated replication in the presence of novobiocin (55% and 66% under 5x IC_50_ and 10x IC_50_, respectively; magenta arrows). In the latter group, 81% of the cells in the 5xIC_50_ group (n=40) but only 17% of the cells in the 10x IC_50_ group (n=20) initiated the next replication round during antibiotic treatment. In those cells, we observed significant prolongation of the B+D period (86 +/− 74 min and 101 +/− 64 min under 5x IC_50_ and 10x IC_50_, respectively). Interestingly, we found that the proportion of cells that underwent multifork replication was larger under 5x IC_50_ novobiocin than under the optimal conditions (16% vs. 11%, respectively), whereas this fraction dropped to 3% under 10x IC_50_ novobiocin. This is likely to reflect the lengthening of the C period, and suggests that multifork replication may be a key strategy for bacterial survival under non-optimal conditions. The decrease of this fraction under 10x IC_50_ novobiocin may suggest that even though multifork replication serves as an initial response to replicative stress, higher concentrations of novobiocin lead to further gyrase inhibition and may affect global organization of the chromosome to prevent initiation (and re-initiation) of subsequent rounds of replication. In cells undergoing replication under novobiocin treatment, we clearly observed splitting of ParB-mCherry foci, suggesting that the segregation of nascent *oriCs* was not significantly impaired; however, the ParB foci were closer together within cells exposed to novobiocin (see kymograph in Fig. 2B). During novobiocin exposure, cells elongated only 30% (5x IC_50_) to 50% slower (10x IC_50_) than seen under the optimal conditions. Interestingly, after removal of novobiocin, we frequently observed additional spots of DnaN-mCherry that did not colocalize with α-YFP, indicating that the beta clamp is involved in processes other than chromosome replication (e.g., DNA repair) and/or that residual sliding clamps accumulate on the lagging DNA strand as shown previously (56–59).

**Fig. 2.**
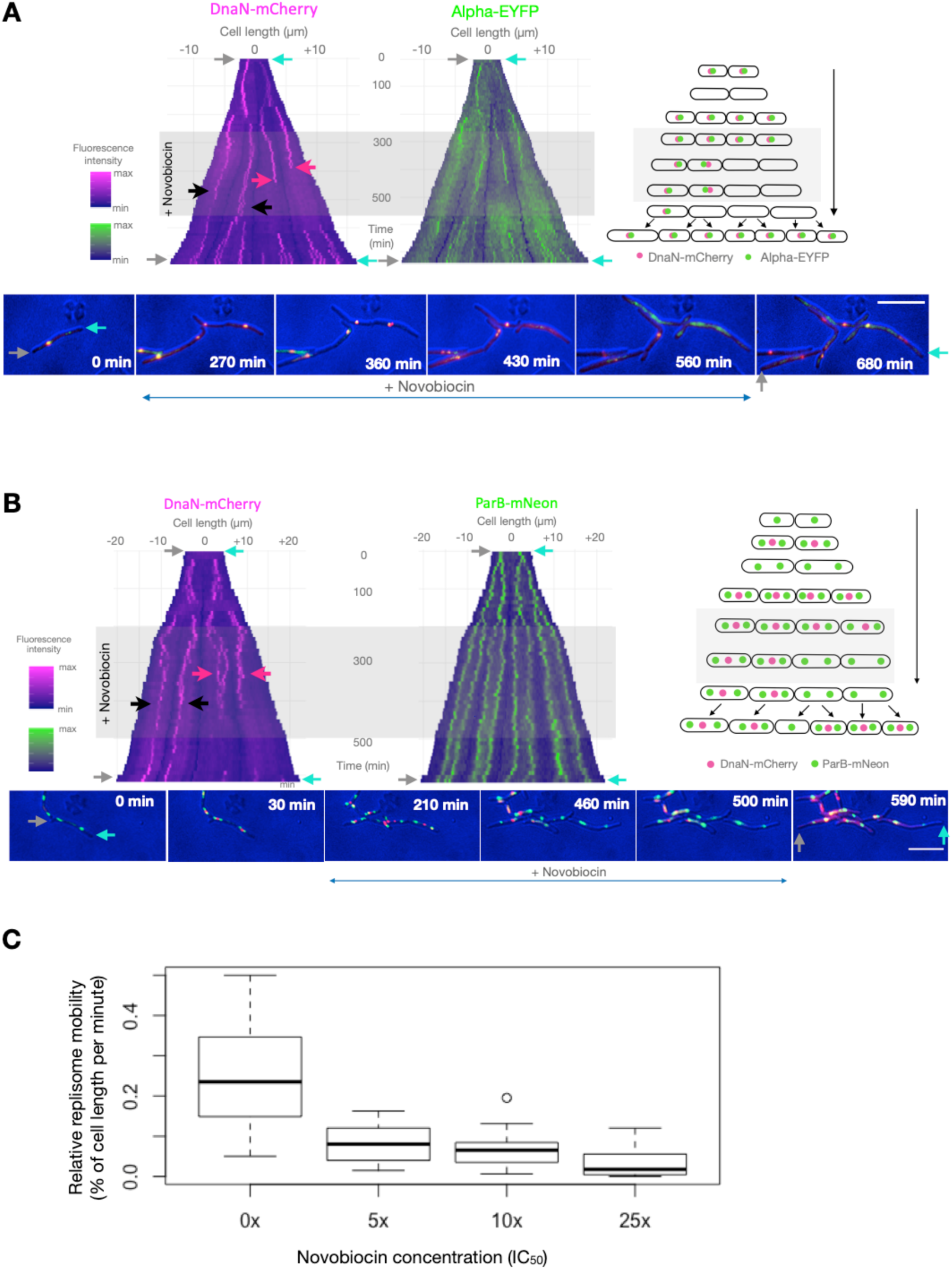
Novobiocin stalls replisomes but has only a moderate effect on growth in *M. smegmatis* cells. Kymographs of representative cells of two *M. smegmatis* strains exposed to novobiocin: DnaN-mCherry/Alpha-EYFP (A) and DnaN-mCherry/ParB-mNeon (B). The left kymographs show DnaN-mCherry fluorescence over time (A and B), while the right ones present Alpha-EYFP (A) or ParB-mNeon (B) fluorescence over time. Time-lapse images of the cells depicted on the kymographs are presented in the bottom panel and a schematic graph of analyzed cells is presented in the right panel. Antibiotic treatment is indicated by gray rectangles; black arrows mark cells in which replication proceeds in the presence of novobiocin; red arrows mark cells in which replication terminates during novobiocin treatment; gray and cyan arrows indicate the corresponding poles on both images and kymographs. Scale bar, 5 µm. (C) Dose-dependent inhibition of replisome mobility during novobiocin exposure of DnaN-mCherry/ParB-mNeon cells. Box-plots represent replisome mobility (defined as the relative distance a replisome traveled along the cell per minute) under increasing concentrations of novobiocin (0x, 5x, 10x and 25x IC_50_, n=20 for each concentration). Measurements were performed from the old pole.

### Addition of nalidixic acid results in growth arrest and replisome disassembly

In contrast to novobiocin, the addition of Ndx (at both 10x IC_50_ and 5x IC_50_) resulted in replisome collapse (Fig. 3A and B, Supp. Movie 2). The timing of replisome disassembly was proportional to the applied concentration of Ndx, occurring at 40 min for 10x IC_50_ but 214 min for 5x IC_50_ (n=45 and 50 cells, respectively). As a consequence of the replisome collapse, the *oriC*s failed to properly segregate. This is intuitive, as after replisome disassembly newly replicated regions do not emerge behind the forks and that stops the segregation process. As shown in Figure 3B, in cells that had begun replicating shortly before being switched to Ndx-containing medium, the ParB complexes remained in close proximity rather than reaching their native positions proximal to the cell poles. This pole-proximal localization of ParB-mNeon was not restored in these cells until very late during the washout period. Unexpectedly, after Ndx washout, replisomes assembled at the same site from which they had previously disassembled in approximately 70% of cells in both the DnaN-mCherry/ParB-mNeon and DnaN-mCherry/α-YFP strains (n=100 per strain).

**Fig. 3.**
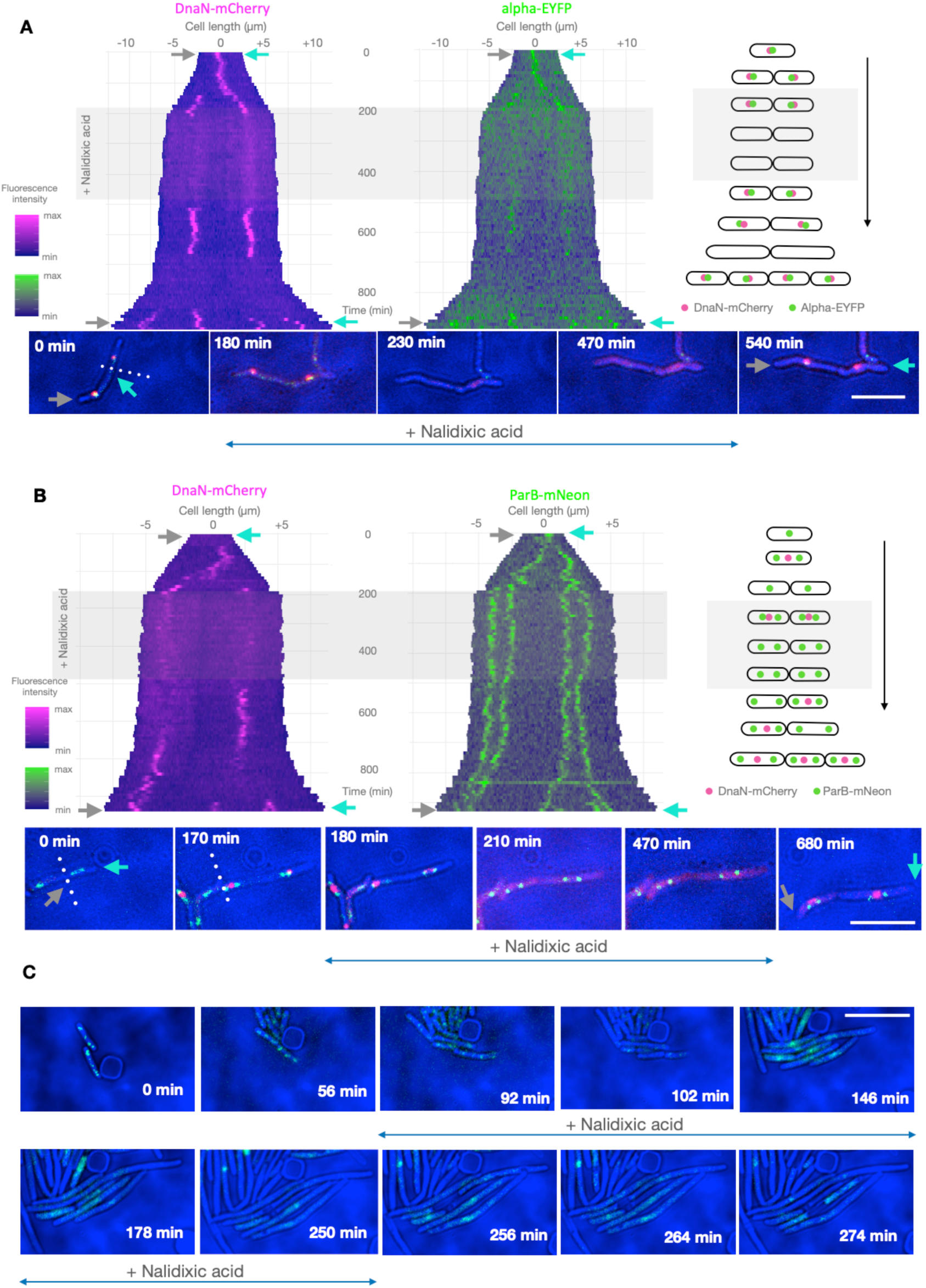
Nalidixic acid treatment results in replisome disassembly and growth arrest of *M. smegmatis* cells. Kymographs of representative cells of two *M. smegmatis* strains exposed to Ndx: DnaN-mCherry/Alpha-EYFP (A) and DnaN-mCherry/ParB-mNeon (B). The left kymographs show the DnaN-mCherry fluorescence over time, while the right ones present the Alpha-EYFP (A) or ParB-mNeon (B) fluorescence over time. Time-lapse images of the cells depicted on the kymographs are presented in the bottom panel, and the schematic graph of analyzed cells is presented in the right panel. Antibiotic treatment is indicated by the gray rectangle; gray and cyan arrows indicate the corresponding poles on both images and kymographs; and the dotted line indicates the boundary between two daughter cells. (C) TLMM analysis of *E. coli* cells harboring the YPet-DnaN fusion, as assessed during Ndx treatment. Scale bar, 5 µm.

*M. smegmatis* cells stopped growing soon after the addition of Ndx, and the timing of growth restoration during the washout period was not dependent on the utilized concentration of the drug. This growth arrest in *M. smegmatis* was strikingly different from our results obtained in *E. coli* (YPet-DnaN strain kindly provided by dr. Reyes-Lamothe (60)) subjected to a similar Ndx treatment scheme (the exception was the use of a 2-hour Ndx exposure which is also twice the duration of the C period in this bacterium (61)). Consistent with previous reports (62, 63), we observed that *E. coli* cells became filamentous during Ndx treatment, indicating that there was dysregulation between cell growth and division. Additionally, in our study a single YPet-DnaN focus appeared transiently in the central part of each filamentous *E. coli* cell (Fig. 3C). It remains unclear whether the YPet-DnaN foci observed in Ndx-treated *E. coli* cells are due to the replication hold-up rather than the involvement of the beta clamp in the DNA repair of double-strand breaks in the chromosome. Interestingly, elongating Ndx-exposed *E. coli* cells observed in the DIC channel exhibited the transient formation of “holes” that presumably reflected differences in the density of the cytoplasm. This was not accompanied by any disruption of the cell integrity, as all cells were viable and their growth was not arrested. Similar observations were also described in previous studies (62), in which the holes were regarded as vacuole-like structures. The filamentation of *E. coli* cells was attributed to initiation of the SOS response after the induction of double-strand breaks by Ndx (64, 65). In the case of *M. smegmatis*, in contrast, we did not observe similar changes in Ndx-exposed cells, nor did we observe any of the additional DnaN spots observed in the corresponding novobiocin-treated cells.

### Griselimycin affects replisome processivity and leads to the formation of *oriC*-proximal loops

Unlike novobiocin and Ndx, which impose indirect effects on replisomes, griselimycin (GM) potently inhibits the interaction of DnaN (a beta clamp) with the catalytic subunit alpha in the core of DNA Pol III (Fig. 1A). Thus, the Alpha-EYFP/DnaN-mCherry strain was ideal for investigating the action of GM, as the interacting proteins are tagged with different fluorophores in this strain. Based on previous report (43), we expected GM to block the proper assembly of replisomes *in vivo*. Indeed, after GM was added to *M. smegmatis* cells, the DnaN-mCherry foci rapidly disappeared and the fluorescence signal remained diffuse for the rest of the antibiotic treatment period (Fig. 4A-C, Supp. Movie 3). Strikingly, we also observed several appearances of short-lasting Alpha-EYFP fluorescent foci during GM exposure; the lifetimes of these foci were much shorter than the C period in the absence of antibiotic treatment, suggesting that cells presenting such foci underwent abortive replication or replication restart after replisome collapse. To investigate this further, we used an additional reporter strain that expressed Alpha-EYFP and ParB-mCherry from their native chromosomal loci. TLMM analysis revealed that during GM treatment, the appearance of the Alpha-EYFP foci was accompanied by ParB-mCherry (*oriC*) duplication, suggesting that a new round of replication had been initiated. The Alpha-EYFP foci disappeared soon after duplication of the ParB-mCherry complex (39 +/− 20 min in the 10x IC_50_ group, as assessed at 2-min frame intervals; n=47). During GM treatment, we observed up to several duplication events at the *oriC* region, each of which was preceded by the colocalization of Alpha-EYFP with ParB-mCherry. To emphasize, we did not observe colocalization of DnaN-mCherry with Alpha-YFP foci in DnaN-mCherry/Alpha-YFP cells exposed to GM. Hence, during GM exposure, the replisome can assemble in the *oriC* region, but its inability to interact with the sliding clamp results in loss of processivity of the core polymerase and consequent abortive replication. Several loops containing an *oriC*-proximal region might therefore arise (which is consistent with the presence of multiple ParB-mCherry foci in a single cell, Fig. 4C) and the length of these loops presumably correlates with the lifetime of individual Alpha-EYFP foci.

**Fig. 4.**
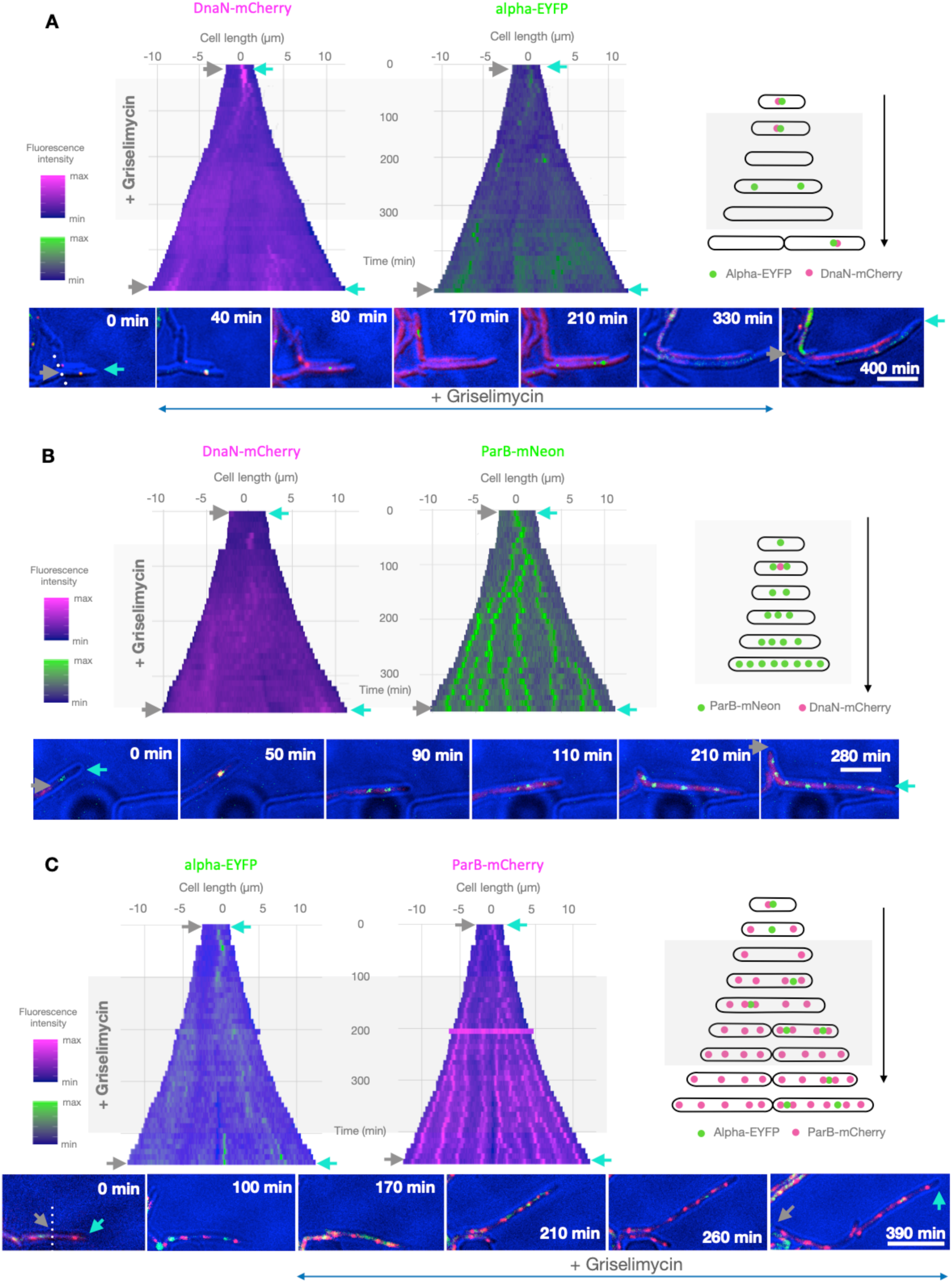
Griselimycin treatment results in loss of replisome processivity and multiplication of *oriC*-proximal regions. Kymographs of representative cells of three *M. smegmatis* strains exposed to GM: DnaN-mCherry/Alpha-EYFP (A), DnaN-mCherry/ParB-mNeon (B) and Alpha-EYFP/ParB-mCherry (C). The left kymographs show the fluorescence of DnaN-mCherry (A, B) or Alpha-EYFP (C) over time, while the right ones present the fluorescence of Alpha-EYFP (A), ParB-mNeon (B) or ParB-mCherry (C) over time. Time-lapse images of the cells depicted on the kymographs are presented in the bottom panel and a schematic graph of analyzed cells is presented in the right panel. Antibiotic treatment is indicated by the gray rectangle; gray and cyan arrows indicate the corresponding poles on both images and kymographs; and a dotted line indicates the boundary between two daughter cells. Scale bar, 5 µm.

Interestingly, the fluorescence intensities of both Alpha-EYFP and DnaN-mCherry varied significantly across individual cells during GM washout, but not during the pre-treatment period. This variation in fluorescence intensity was only partially due to replisome assembly, as we observed high-intensity dispersed fluorescence in addition to well-defined foci. This was particularly noticeable in the DnaN-mCherry/Alpha-EYFP strain. During washout, we observed different levels of DnaN-mCherry and Alpha-EYFP in various cells even within a single microcolony (Fig. S3). This clearly supports the idea that bacterial heterogeneity can arise under stressful conditions (e.g., under antibiotic treatment) and may be a key factor not only for survival under stress but also during the recovery phase and the recolonization of the previously occupied niche. In addition to the heterogeneity of fluorescence intensities, we also observed that the DnaN-mCherry foci were generally overrepresented in comparison to Alpha-EYFP during the washout period. Only some of the DnaN-mCherry complexes colocalized with Alpha-EYFP, confirming that the beta clamp could be involved in DNA repair and/or recombination events.

### Chromosome dynamics are differentially affected by the various replication inhibitors

In addition to its effects on transcription, DNA replication influences the overall nucleoid organization (66–70). We thus examined how antibiotic-triggered replisome collapse (Ndx), replication fork hold-up (novobiocin) or the loss of processivity (GM) affected the nucleoid structure. To answer these questions, we used the DnaN-mCherry/HupB-EGFP strain, which allowed us to simultaneously observe chromosome dynamics (HupB is a homolog of the HU protein from *E. coli*, which occupies the whole nucleoid and is used as a chromosomal marker (53, 54, 71)) and track the progression of replication. Our previous studies demonstrated that mycobacterial nucleoid adopts a bead-like structure spread along the cell. However, positioning of the nucleoid is asymmetric, being closer to the new cell pole for almost the entire cell cycle (54). Antibiotic exposure of DnaN-mCherry/HupB-EGFP cells yielded replication patterns similar to those described in the other strains of *M. smegmatis* exposed to each drug (see above). Interestingly, we observed that the various antibiotic-induced alterations in replisomes dynamics reflected different changes in chromosome organization.

The area occupied by the mycobacterial chromosome, which was measured by comparing the total area occupied by HupB-EGFP before and during antibiotic treatment, was decreased by exposure to nalidixic acid (10x IC_50_) (Fig. 5A, B and E). The chromosome shrank rapidly after Ndx was introduced to the medium (29 +/− 7.5 min, n=57), then decondensed slowly during washout, beginning at 120 min after the removal of Ndx. The chromosome was condensed to approximately 30% of its initial size (27 +/− 20%, n=49), which accounts for the observed increase in the fluorescence intensity of HupB-EGFP. Interestingly, in cells that initiated replication less than 60 minutes before being switched to Ndx-containing medium (i.e., those in which replication had progressed less than halfway) the chromosome area shrank to the point that it was visualized as a single fluorescent cluster per cell (Fig. 5A and the left cell in 5E). In contrast, in the cells in which replication had proceeded more than halfway before the addition of the drug (i.e., those that initiated replication a minimum of 60 min before Ndx addition), the chromosome was compacted to the point that we observed two separate fluorescent clusters (Fig. 5B and the right cell in 5E). These clusters presumably reflected newly replicated sister chromosome regions. In this, our observations parallel the previous description of bilobed chromosomes in slow-growing *E. coli* cells (72, 73). The characteristic bead-like structure of the mycobacterial chromosome was lost upon Ndx treatment, to be replaced by a large uniform fluorescent patch. Chromosome compaction and replisome disassembly together explain why the nascent *oriC*s remained in close proximity throughout Ndx treatment and even longer after its removal. The changes in nucleoid density were reversible in all observed cells, and the chromosome regained its normal morphology (i.e., a bead-like pattern) after Ndx washout. Our observations are in line with those previously obtained in Ndx-exposed *E. coli* cells (62, 63), which showed that the chromosome was compacted around the midcell. However, we did not observe decondensation of the chromosome during prolonged Ndx exposure, as previously reported in *E. coli* (62). We hypothesize that the decondensation seen in Ndx-treated *E. coli* reflected fragmentation subsequent to the introduction of double-strand breaks. Our present results suggest that, unlike *E. coli* cells, *M. smegmatis* cells do not undergo chromosome fragmentation upon Ndx treatment.

**Fig. 5.**
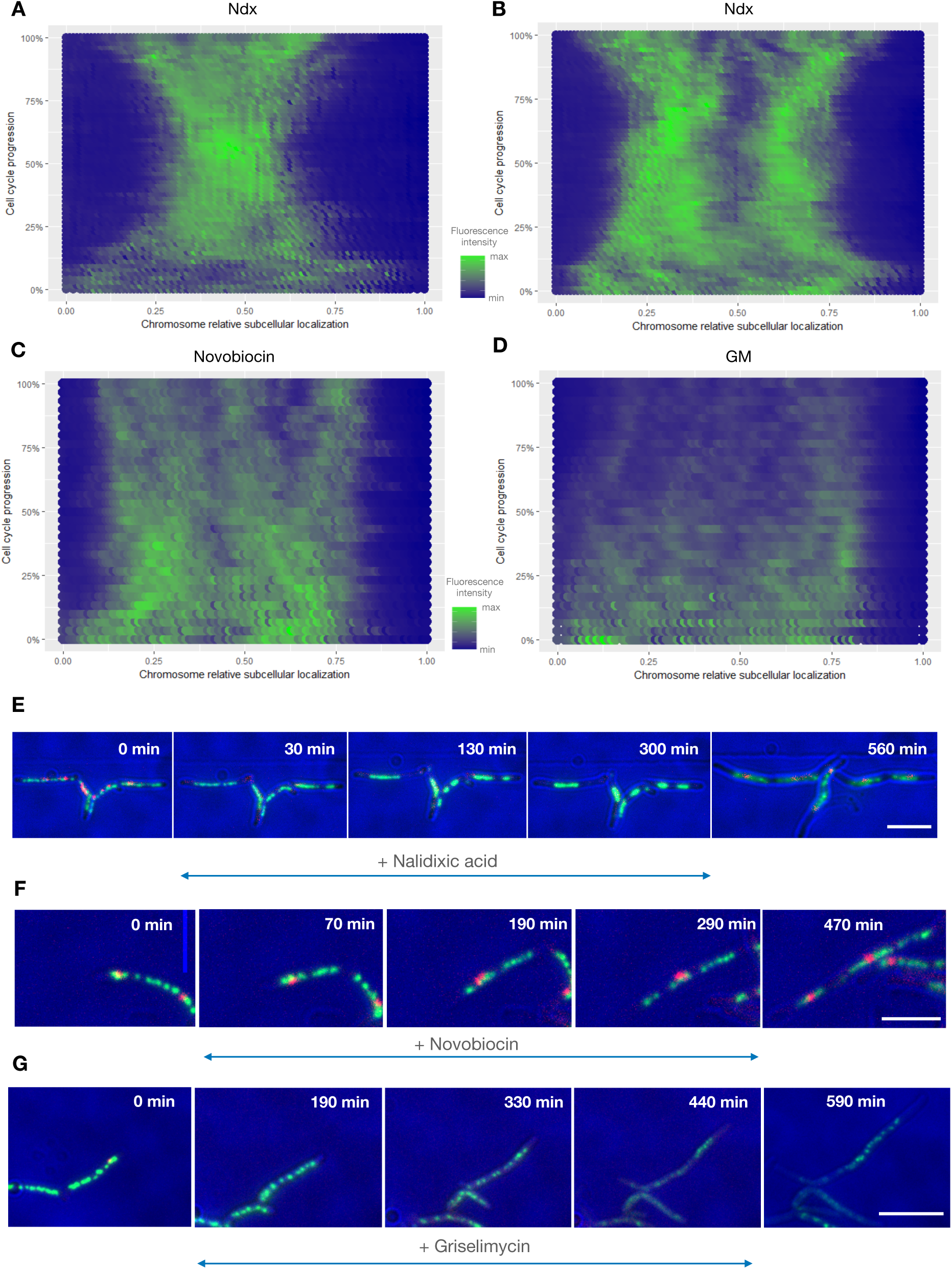
All three analyzed antibiotics affect chromosome organization and dynamics. A-D) Fluorescence profiles of HupB-EGFP over time for representative cells (n=5) under treatment with the different antibiotics. (A) In cells with overall replication progression of <50%, (cells in which replication progressed beyond the halfway point). Ndx exposure causes the chromosome to shrink until only a single fluorescent cluster is visible. (B) In cells with overall replication progression of >50%, Ndx exposure causes the chromosome to shrink until two fluorescent clusters are visible. (C) During novobiocin treatment, chromosome decondensation is visualized as separate fluorescent clusters. (D) During GM treatment, chromosome decondensation is followed by a decrease in the fluorescence intensity. (E-G) TLMM images of representative cells under treatment with Ndx (E), novobiocin (F) or GM (G). Scale bar, 5 µm.

In contrast to the effects observed in Ndx-treated cells, those exposed to novobiocin exhibited preservation of the bead-like chromosome structure throughout the antibiotic treatment (Fig. 5C and F). Even in the 10x IC_50_ group, replication still proceeded (albeit with a delay), as did cell elongation. We observed chromosome decondensation, which presumably reflected the increased cell volume. The area occupied by the nucleoid was extended, but the intensities of the HupB-EGFP foci were decreased only marginally. Notably, the chromosomes of novobiocin-treated cells formed clusters consisting of regular HupB-EGFP foci, which were further apart along the long cell axis compared to the foci of actively replicating cells growing in novobiocin-free medium. This may indicate that newly replicated sister chromosomes are more likely to separate in filamentous cells.

In cells exposed to GM, the chromosome structure underwent dynamic rearrangements. As seen for novobiocin treatment, GM treatment triggered chromosome decondensation; however, the area occupied by the nucleoid was much larger under GM exposure than under novobiocin exposure (Fig. 5D and G). In most GM-treated cells, the HupB-EGFP foci were heterogeneous in size and intensity, and they were distributed unevenly along the cell (Fig. 5G, top cell). Moreover, in 17% of the cells (n=155 cells), the bead-like pattern of the HupB-EGFP complexes was lost and only dispersed fluorescence was observed (Fig. 5G, bottom cell). We hypothesized that this might reflect that HupB binding is diminished by the over-relaxation of the chromosome, which fills more space in growing filamentous cells, and/or the induction of DNA damage, as seen in Ndx-treated *E. coli* strains (62). To test the first possibility, we treated cells with GM for 5 hours and performed Hoechst33342 staining. This staining overlapped with the fluorescence arising from the HupB-EGFP complexes (Fig. S4), indicating that there was no change in the binding of HupB to the chromosome of GM-treated *M. smegmatis* cells. This suggests that the dispersed HupB-EGFP fluorescence seen in some cells is likely to reflect a change in chromosome organization. Indeed, further analyses revealed that 50% of cells that exhibited dispersed fluorescence (n=20) continued to grow and 37% of them regained the normal chromosome structure after GM washout. The remaining 50% of cells did not elongate further and were believed to be dead; notably, their dispersed fluorescence signal was preceded by the sudden condensation of the nucleoid to a single bright HupB-EGFP spot, which then rapidly disappeared. Thus, the decrease of the HupB-EGFP intensity and loss of regular chromosomal pattern may contribute to the DNA damage, at least in some of the cells.

In summary, our present results indicate that treatment of *M. smegmatis* cells with Ndx, novobiocin or griselimycin induced various rearrangements of the chromosome structure, reflecting differences in the action modes of the tested antibiotics.

## Discussion

Here, we present a novel system that allows to monitor in real time the action of antibiotics at the single-cell level. Using time-lapse microfluidic microscopy (TLMM) and a set of reporter strains, we show how replication-affecting drugs alter the replisome and chromosome dynamics of mycobacterial cells. To date, only a few studies have examined the action of antimicrobials at a single-cell level in real time. Given that antibiotic treatment increases bacterial heterogeneity (41, 42), such studies are needed to shed light on bacterial physiology during antibiotic exposure and after its removal. Our present work reveals that combining single-cell techniques with appropriate target-tagged strains can provide new insights into the action mechanisms of the tested antimicrobial compounds. We chose *M. smegmatis* as a well-established model for studying the biology of the tubercle bacilli, some of which (e.g., *M. tuberculosis*, *M. bovis*) cause severe diseases that have enormous impacts on global health and the world economy. The use of such a model is important because mycobacteria substantially differ from the extensively examined model bacteria, such as *E. coli*, *B. subtilis* and *C. crescentus*. In particular, asymmetric septum placement and variations in the elongation rates at the cell poles of mycobacteria create heterogeneous populations in which individual cells may respond differently to stress-inducing stimuli (30, 31, 39).

We tested three replication-affecting drugs (Fig. 1): Nalidixic acid (Ndx) and novobiocin target different sites of the heterotetrameric DNA gyrase (GyrA and GyrA/GyrB/DNA, respectively) (44, 48) to alter replisome passage, whereas griselimycin (GM) prevents the beta clamp from interacting with the core DNA polymerase III complex, causing the replisome to lose its processivity. The tested inhibitors had various impacts on the dynamics of the replication complex. Novobiocin stalled the replisomes (observed as prolongation of the C period and a decrease in the subcellular mobility of the tagged replisomes) and moderately decreased the cell elongation rate (30% and 50% decreases under 5x IC_50_ and 10x IC_50_, respectively). In contrast, Ndx completely halted replication, as indicated by the disappearance of the replisome foci. Surprisingly, unlike the situation in *E. coli* (Fig. 3C and (62, 63)), Ndx arrested the growth of mycobacterial cells. This may suggest that the action mechanism of Ndx may differ between bacteria. Finally, we found that GM aborted the interaction of the core DNA polymerase III complex with the sliding clamp, which was observed as a loss of colocalization for DnaN-mCherry and Alpha-EYFP. Interestingly, Alpha-EYFP foci (but not DnaN-mCherry foci) were still observed during GM treatment, but their lifetime was profoundly shorter than the C period in WT *M. smegmatis*. Additionally, the appearance of Alpha-EYFP foci during GM exposure was followed by duplication of the ParB-mCherry focus, which was attributed to duplication of the *oriC* region. These results suggest that the core polymerase can assemble at *oriC*, but it shows poor processivity due to the loss of the beta clamp interaction. This is in line with previous reports showing that the hydrophobic cleft through which GM binds to the beta clamp contributes to both catalytic subunit interactions and the binding of the clamp loader to the delta subunit (59). Blockage of the hydrophobic cleft by GM probably abolishes the loading of the beta subunit, but not the Alpha subunit (which binds with the tau subunit), onto the DNA strand. As a result, replication is initiated in the presence of GM, but loss of processivity of the replicative complex, probably mediated through a dynamic turnover of replisome subunits (particularly of the core complex) (74, 75), leads to the generation of chromosomal loops consisting of newly replicated DNA fragments. If we combine the mean lifetime observed herein for the Alpha-EYFP focus during GM treatment (39 +/− 20 min) with the calculated DNA synthesis velocity in Alpha-EYFP strains giving 390 bp/sec and assume that this rate is not decreased by the loss of the interaction with the sliding clamp, we predict that the generated loops will be 1.85 Mbp in length (mean; range 565 kbp to 3.9 Mbp; n=47) and cover the *oriC*-flanking chromosomal fragment. We hypothesize that these loops may undergo recombination, which in turn may cause multiplication of *oriC*-proximal regions. In fact, multiplication of the fragment encompassing *oriC* (and the *dnaN* gene) has been reported in GM-resistant strains of *M. smegmatis* (43). During the GM washout period, we observed multiple DnaN-mCherry foci, only some of which colocalized with the Alpha-EYFP complexes. We speculate that these DnaN-mCherry foci may be involved in aforementioned recombination between the *oriC*-proximal loops.

All three tested antibiotics also impacted the overall nucleoid structure. Novobiocin displayed the most modest effect. The bead-like structure of the chromosome was maintained throughout novobiocin treatment, although there was an increase in the distance between individual nucleoid clusters. This stretching of the area occupied by the nucleoid was probably a consequence of the increased cell volume of filamentous cells. GM treatment triggered a more pronounced decondensation of the nucleoid: the bead-like pattern was lost and there was a decrease in the fluorescence intensity of HupB-EGFP. Loss of the bead-like structure might reflect either over-relaxation of the chromosome or chromosome fragmentation and subsequent cell death (similar to that observed in quinolone-treated *E. coli*; (62)). Because the genes encoding both subunits of DNA gyrase are located very near *oriC* in *M. smegmatis* (approx. 5 kb away from the *oriC*), the generation of *oriC*-proximal loops might result in multiplication of the *gyrA* and *gyrB* genes and the consequential increased level of gyrase. This might, in turn, affect global supercoiling and overall chromosome organization. This hypothesis is supported by our observation that the Ndx-mediated blockade of gyrase activity led to chromosome compaction. Temporal loss of the regular chromosome structure was also observed in a fraction of GM-treated cells during the washout period, mirroring the heterogeneity arose during the adaptation to new growth conditions (i.e., the resumption of growth after antibiotic treatment). Interestingly, in some cells exposed to GM, the HupB-EGFP signal was reduced to a single fluorescent spot that became visible for a short time and then disappeared. This was accompanied by a sudden growth arrest that was not reversed during the washout period. In such cells, only dispersed fluorescence was visible. Similar changes were described earlier in *E. coli* cells exposed to quinolones (62, 76). It is possible that frequent collapse of replisomes may, at least in some cells, trigger the SOS response, and that the dramatic decrease in fluorescence intensity may be a hallmark of chromosome fragmentation. Unexpectedly, we did not observe similar changes in Ndx-exposed *M. smegmatis* cells, and the observed chromosome condensation was reversed upon washout in these cells. This phenomenon, which was accompanied growth arrest, was strikingly different from that previously reported in *E. coli* (62, 63). However, it is consistent with previous findings that classical quinolones (including Ndx) do not induce the cleavage activity of *M. tuberculosis* gyrase (77, 78). This is presumably due to the presence of alanine in the 90th position of the GyrA sequence; this corresponds to the conserved Ser83 found in many other bacterial species, which has been implicated (together with an acidic residue four positions downstream) in the ability of GyrA to interact with quinolones via a water-metal ion bridge. The substitution of serine with alanine in the corresponding region of *M. tuberculosis* GyrA is believed to be a key factor responsible for the intrinsic resistance of mycobacteria to quinolones (at least the classical ones). In the present study, we confirmed on the single-cell level that Ndx does not induce double-strand breaks in *M. smegmatis* (the chromosome was condensed rather than fragmented, as was observed for *E. coli*; (62)), and probably therefore, does not induce the mechanism(s) responsible for the SOS response. As mentioned above, gyrase activity-altering antibiotics may affect global supercoiling and thereby alter the expression levels of chromosome topology-regulated genes. In fact, chromosome topology serves as a specific stress-sensing system that enables the expression levels of many genes to be rapidly and simultaneously altered. Importantly, many genes involved in virulence are regulated by changes in DNA topology (79–81).

In addition to determining the single-cell action of replication-affecting drugs, our system also enabled us to observe a high degree of cells heterogeneity.,during both the antibiotic exposure and the washout period. The between-cell differences in replisome and chromosome dynamics, which were more pronounced at lower inhibitor concentrations (i.e., 5x IC_50_ and 10x IC_50_), likely reflected between-cell differences in the cell cycle progression. Future studies measuring individual responses in synchronized bacterial cultures are needed to provide more details about this cell heterogeneity.

Our present work may also shed light on the effectiveness of combining different antibiotics in the treatment of mycobacterial infections. Previous studies showed that combination therapy may yield either synergy or antagonism between various antibiotics (82, 83). Synergy was observed most often when the combined drugs acted on the same process (e.g., cell wall synthesis), such as when different *β*-lactams were combined. In contrast, the combination of drugs belonging to different classes (especially DNA and protein synthesis inhibitors) tended to have an antagonistic effect. Interestingly, strong synergy was observed when an aminoglycosides (e.g., gentamycin) was coupled with *β*-lactams. Antagonism has been observed more frequently than synergy in such studies, indicating that extreme caution is required when combining drugs to battle bacterial infections. Our present results suggest that in mycobacteria combining a cell wall synthesis inhibitor with nalidixic acid (or other quinolones) would not be likely to show any additional benefit beyond the use of Ndx alone, as the cells stopped growing immediately after Ndx was introduced to the medium. On the other hand, it seems that *β*-lactams may be useful when combined with griselimycin or novobiocin. Given that the standard treatment for TB consists of combining rifampicin, isoniazid, pyrazinamide and ethambutol for multiple months (84, 85), the system introduced herein may be useful in efforts to determine the optimal drug combination for use against mycobacterial infections and to test new combinations involving second-line compounds, such as fluoroquinolones. In future, it may also prove beneficial in assessing the usefulness of the combination therapy consisting of newly-developed drugs.

## Methods

### Bacterial strains and culture conditions

Allelic replacement of the genes encoding the alpha (*Msmeg_3178*) or beta (*dnaN, Msmeg_0001*) subunits of DNA polymerase III with the *alpha-eyfp* or *dnaN-mCherry* fusion genes, as well as replacement of the *parB* gene with the *parB-mNeon green* or *parB-mCherry* fusion genes, was performed as previously described (51, 52). Replacement of the *hupB* gene with *hupB-egfp* was performed as described by Holowka and coworkers (53, 54). Mycobacterial strains were cultured in 37°C in 7H9 broth or on 7H10 agar (Difco) supplemented with OADC (BD), 0.05% Tween80 and (when needed) proper additives (kanamycin 50 µg/ml, X-Gal, 2% sucrose). Correct allelic replacement and proper incorporation of the integration vectors were confirmed using PCR and Western blotting. Fusion of the functional fluorescent protein was confirmed by semi-native SDS-PAGE and visualized using a Typhoon phosphoimager. Western blotting was performed using standard procedures (86) with polyclonal anti-mCherry, monoclonal anti-GFP (Santa Cruz Biotechnology), and anti-mNeon antibodies (Chromotec).

### Determination of IC_50_

For the growth curve analysis, cells were cultured in a Bioscreen-C instrument (37°C, high speed, 3 days) with or without the tested inhibitors (concentration ranges: GM, 0.1 – 1 µg/ml; Ndx, 30 – 150 µg/ml; Nov, 0.5 – 9 µg/ml). Data were collected every 20 min using a brown filter (600 nm). The growth rate (OD600/min) was estimated by analyzing the slope of the linearly fitted correlation in the exponential growth phase. The percentage of growth inhibition was calculated by comparing the growth rates obtained in the presence of the tested antibiotics to the growth rate obtained without any inhibitor (which was defined as 100%), as described previously (87). The IC_50_ was calculated for each strain from the inhibition curve plotted using the R-package software, and was taken as the concentration of a particular compound that inhibited the cell growth rate by 50%.

### TLMM

TLMM was performed as previously described (51, 87) using B04A plates with an ONIX flow-control system (Merck-Millipore). Cells loaded into the observation chamber were exposed to fresh 7H9/OADC/Tween for 5 hours, 7H9/OADC/Tween/inhibitor for 5 hours and fresh 7H9/OADC/Tween without antibiotic for 7 hours. All experiments were performed under continuous pressure (1.5 psi) at 37°C. Images were recorded in 2- or 10-min intervals using a Delta Vision Elite inverted microscope equipped with a 100 × or 60 × oil immersion objective. Movies were analyzed with the ImageJ Fiji suite.

## Acknowledgements

We thank Dr. Rodrigo Reyes-Lamothe for providing the RRL-19B strain. We are grateful to Agnieszka Strzałka for providing assistance with data analysis using the R statistical programming language. This study was supported by the National Science Center, Poland (MAESTRO grant 2012/04/A/NZ1/00057). The cost of publication was supported by the Wroclaw Centre of Biotechnology under the Leading National Research Centre (KNOW) program, 2014–2018.

---

**Table S1.**
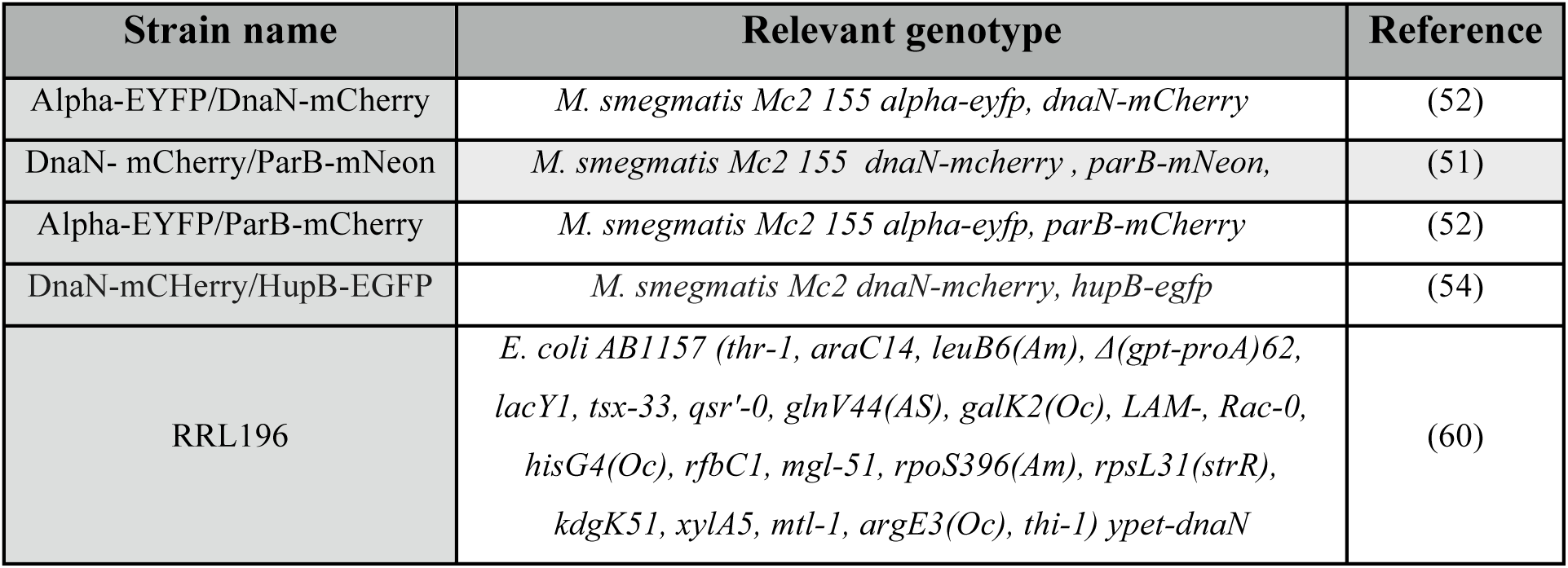
Characterisctics of strains used in this study

**Fig. S1.**
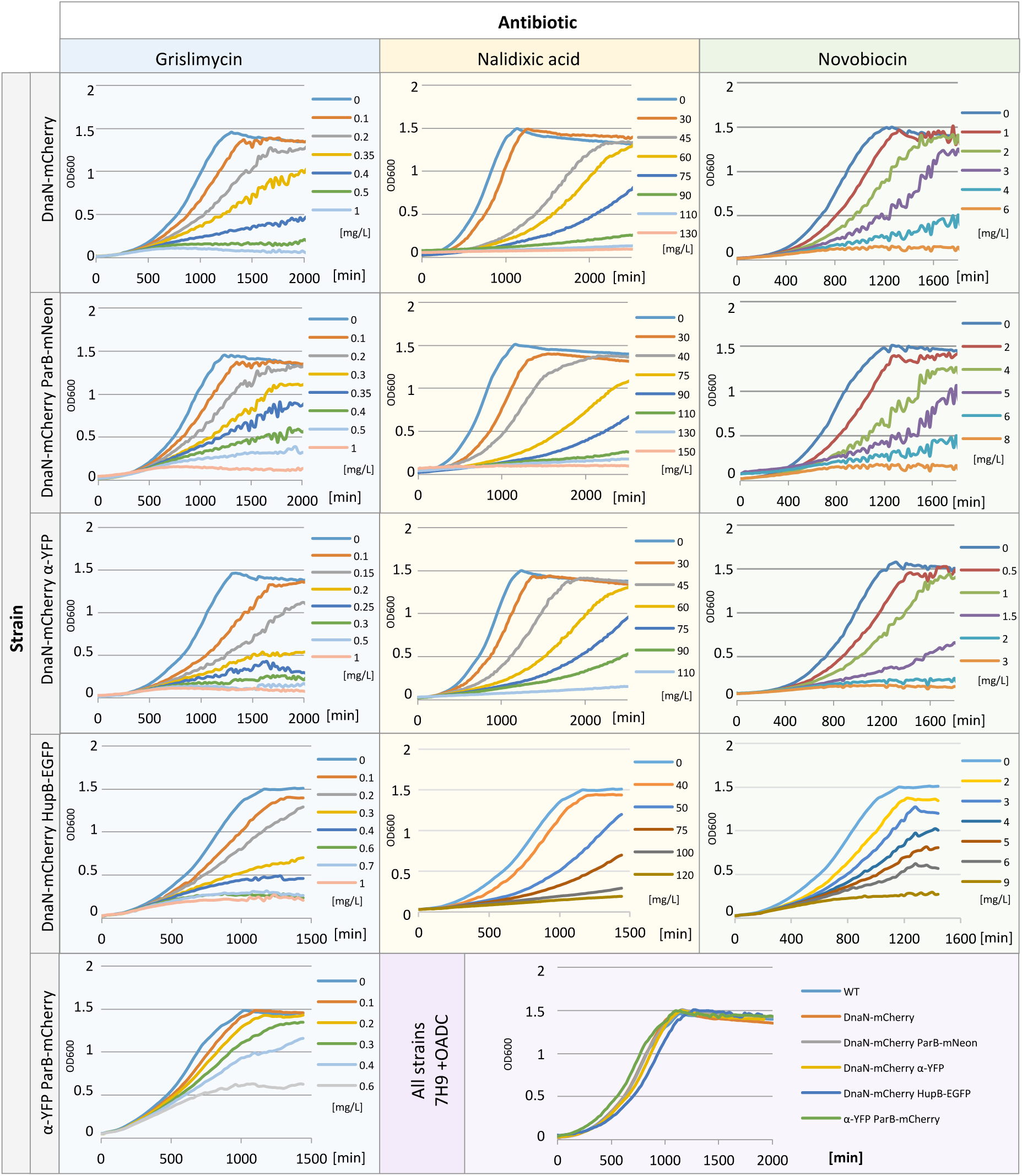
Growth curves of the *M. smegmatis* strains used in this study in the presence of the different antibiotics. The right bottom panel presents the growth of all strains in antibiotic-free medium.

**Fig. S2.**
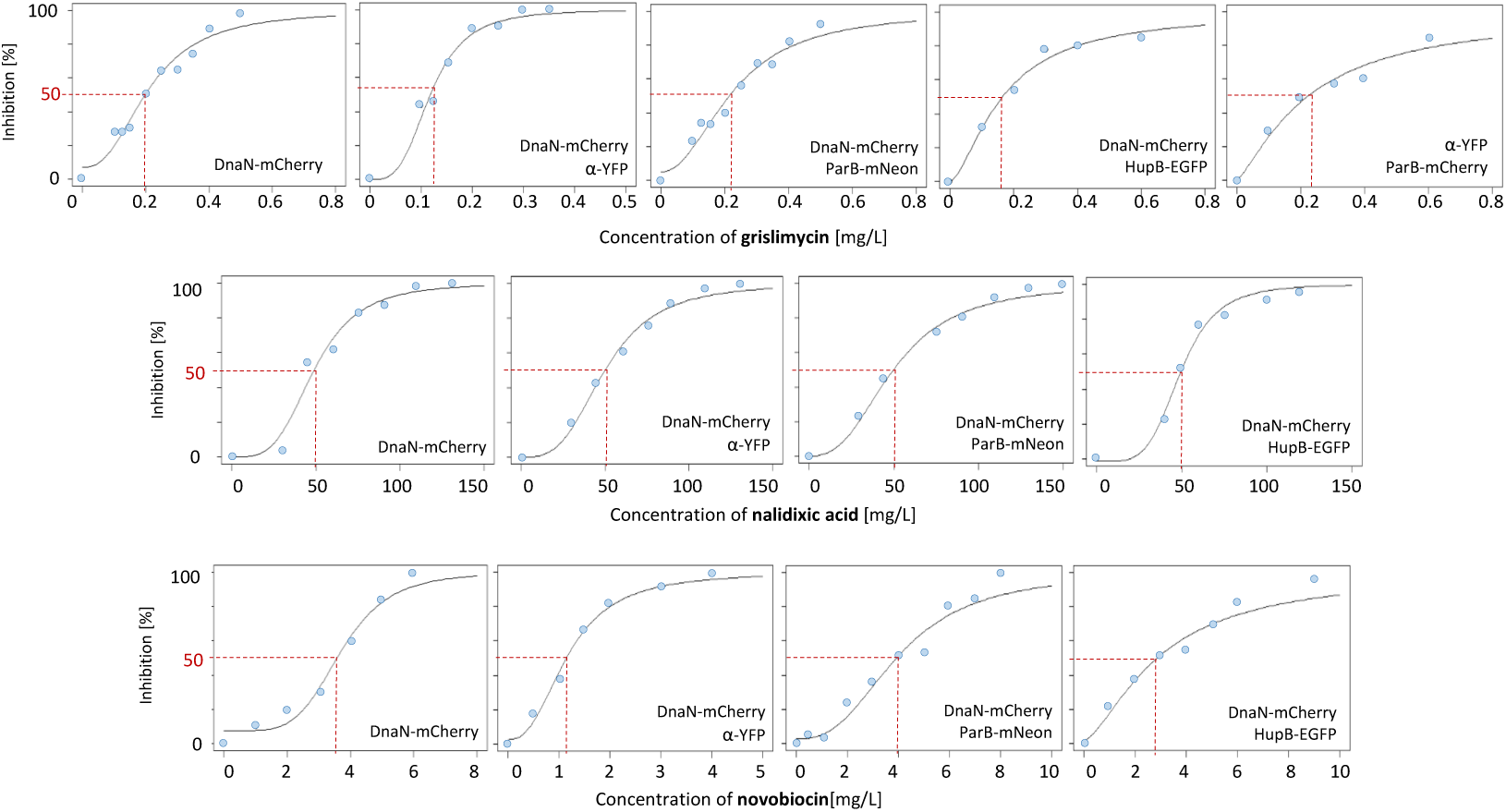
Inhibition curves calculated from the growth curves plotted in Figure S1. Top panel, griselimycin; middle panel, nalidixic acid; bottom panel, novobiocin. IC_50_ is marked with a red dotted line.

**Fig. S3.**
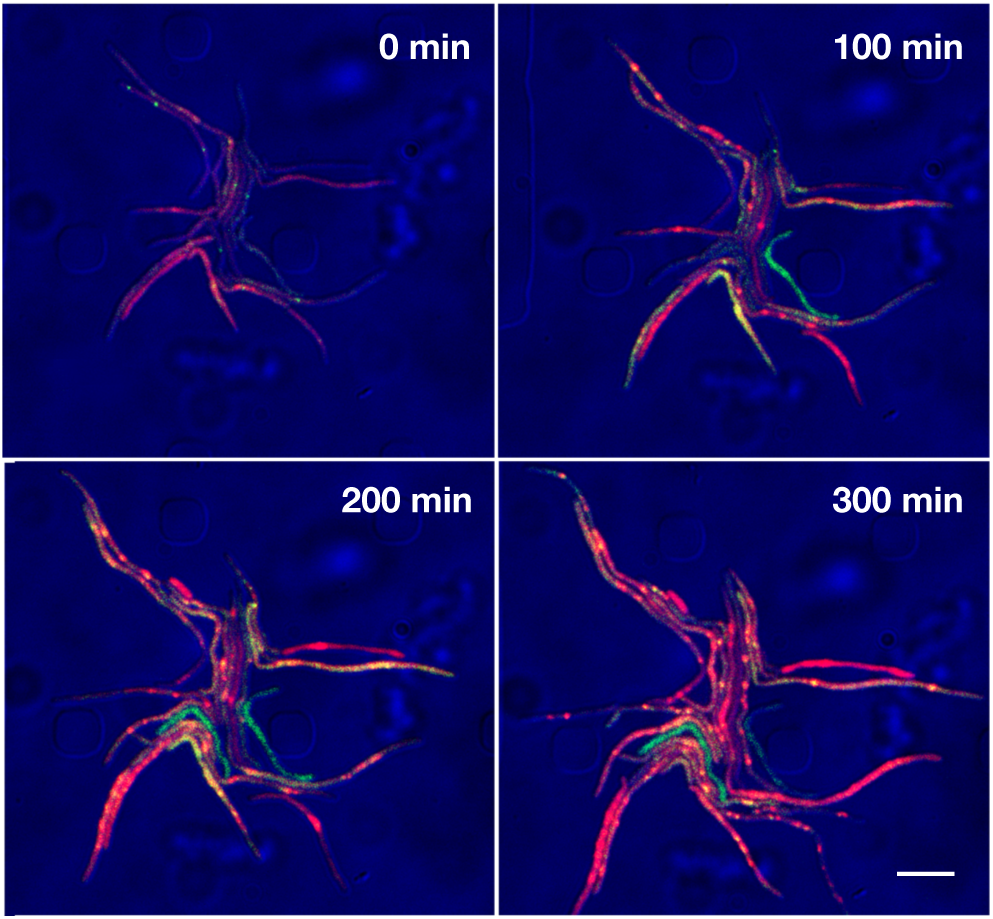
GM treatment induces heterogeneous expression of both Alpha-EYFP and DnaN-mCherry during the subsequent washout period. TLMM images of representative microcolonies of DnaN-mCherry/Alpha-EYFP after the removal of GM. Scale bar, 5 µm.

**Fig. S4.**
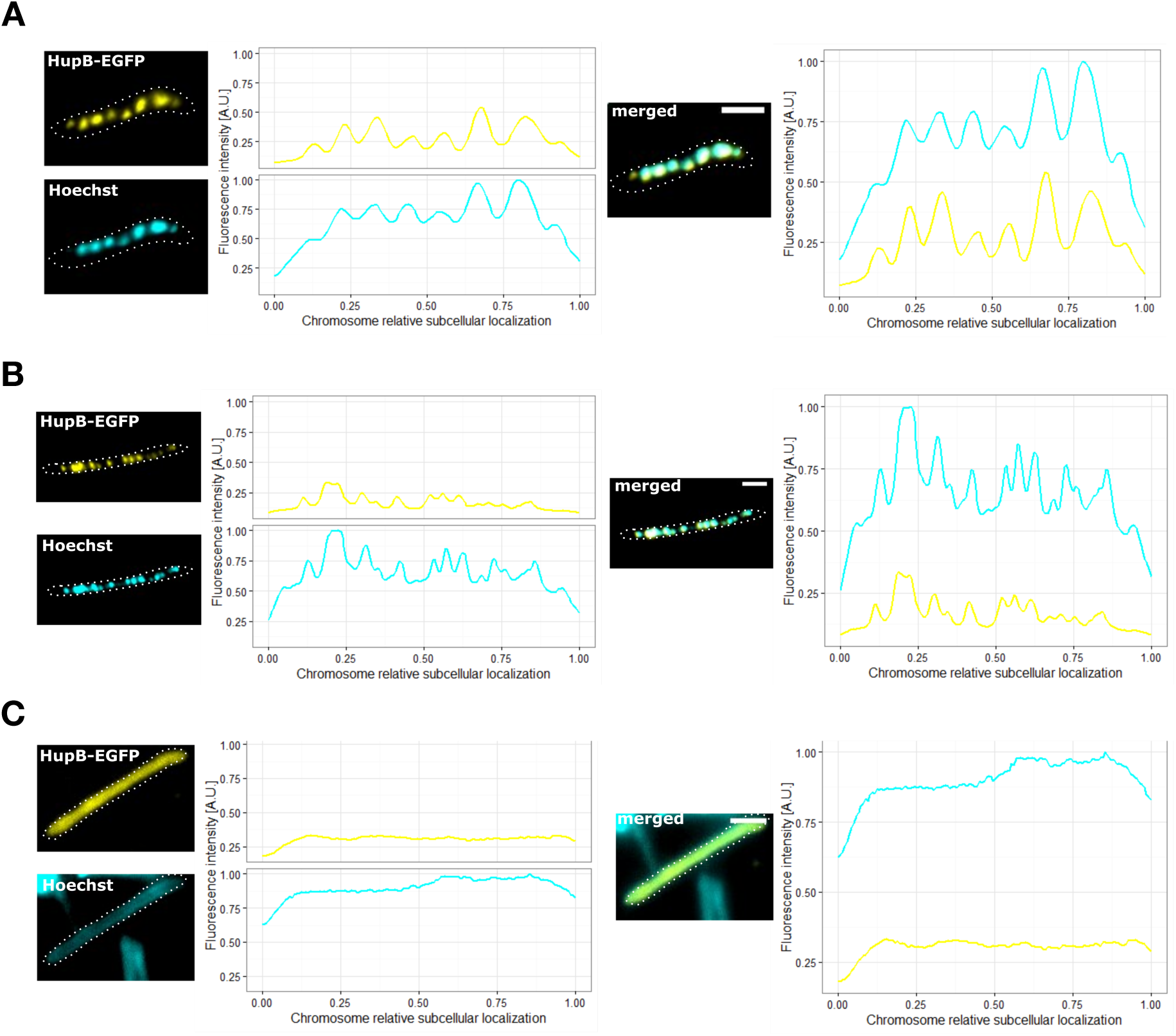
The binding of HupB-EGFP to the chromosome is not affected under GM treatment, and reflects the nucleoid structure. Micrographs and corresponding fluorescence intensity profiles of DnaN-mCherry/HupB-EGFP cells stained with Hoechst 33342 before (A) and after 5 hours of GM treatment (B and C). During GM treatment, the chromosome is decondensed in some cells (B), while diffuse fluorescence is observed in others (C).

